# Roseoflavin, a natural riboflavin analogue, possesses *in vitro* and *in vivo* antiplasmodial activity

**DOI:** 10.1101/2022.04.14.488433

**Authors:** Ayman Hemasa, Matthias Mack, Kevin J. Saliba

## Abstract

The ability of the human malaria parasite *Plasmodium falciparum* to access and utilise vital nutrients is critical to its growth and proliferation. Molecules that interfere with these process could potentially serve as antimalarials. We found that two riboflavin analogues, roseoflavin and 8-aminoriboflavin, inhibit malaria parasite proliferation by targeting riboflavin metabolism and/or the utilisation of the riboflavin metabolites flavin mononucleotide and flavin adenine dinucleotide. An additional eight riboflavin analogues were evaluated, but none were found to be more potent than roseoflavin, nor was their activity on target. Focussing on roseoflavin, we tested its antimalarial activity *in vivo* against *Plasmodium vinckei vinckei* in mice. We found that roseoflavin decreased the parasitemia by 46-fold following a 4 day suppression test and, on average, increased the survival of mice by 4-5 days. Our data are consistent with riboflavin metabolism and/or the utilisation of riboflavin-derived cofactors being viable drug targets for the development of new antimalarials and that roseoflavin could serve as a potential starting point.

## Introduction

Despite constant effort to combat malaria, a disease caused by apicomplexan parasites of the genus *Plasmodium*, the fatality rate remains high, with 627,000 deaths in 2020 (1). Mosquito (the malaria vector) resistance to insecticides and parasite resistance to antimalarials continue to increase in endemic countries (1-3), making malaria control difficult. The danger of acquiring cross-resistance may be increased if new antimalarials are developed which have the same target/s as current antimalarials. In addition, the lack of a highly effective vaccine (4, 5), makes it important to evaluate novel drug targets in order to create safe and effective treatments.

Understanding the essential nutrient requirements of the intraerythrocytic stage of the malaria parasite (the stage responsible for the morbidity and mortality associated with malaria) may shed light on the metabolic pathways that can be used as novel targets. Whilst progress has been made in our understanding of the parasite’s requirement for certain vitamins (e.g. pantothenate (6-8)), very little is known about the parasite’s requirement for other vitamins, such as riboflavin (vitamin B_2,_ see **Figure 1** for structure).

**Figure 1:**
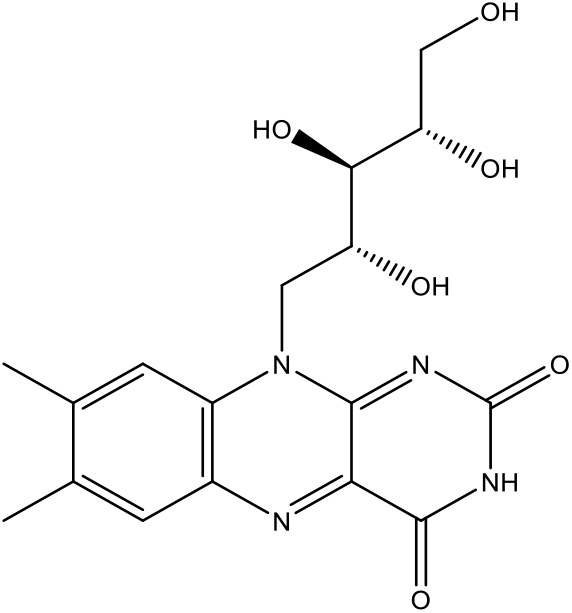
The chemical structure of riboflavin.

Riboflavin is phosphorylated by the enzyme flavokinase into flavin mononucleotide (FMN) which can be adenylylated to flavin adenine dinucleotide (FAD) by the enzyme FAD synthetase. Riboflavin itself has no known biological activity, but its metabolites FMN and FAD (referred to as flavin cofactors) are essential for the activity of flavoenzymes. These enzymes are involved in a variety of biological processes such as redox reactions, electron transport, protein folding, apoptosis, chromatin remodelling, DNA repair, hydrogenation and dehydrogenation processes, and hydroxylation (9, 10). The genes encoding enzymes involved in riboflavin biosynthesis in other organisms (11-16), do not appear to be present in the *P. falciparum* genome (PlasmoDB). Therefore, the host is presumably the source of riboflavin for the malaria parasite. It has been reported that riboflavin uptake and its conversion into FMN and FAD is increased in erythrocytes infected by *P. falciparum* compared to uninfected erythrocytes, consistent with the parasite requiring an extracellular supply of riboflavin (17). In both *Plasmodium lophurea* (18) and *Plasmodium berghei* (19) infections, riboflavin deficiency has been shown to have an inverse relationship with parasitemia. Moreover, in Papua New Guinea, riboflavin deficiency has been found to provide partial protection to newborns infected with malaria (20). However, removing extracellular riboflavin has also been reported to have no effect on parasite proliferation (21), although the erythrocytes may not have been depleted of intracellular flavin stores at the start of that experiment.

A number of riboflavin analogues have shown antibacterial, anticancer, and antiviral activity, specifically by interfering with the metabolism of riboflavin (22, 23). 8-Demethyl-8-methylamino riboflavin was reported to possess *in vitro* activity against *P. falciparum* (21), and 10-(4’-chlorophenyl)-3-methylflavin has been shown to kill *P. falciparum* in culture and *P. vinckei* in mice (24, 25). The antiplasmodial activity of roseoflavin (RoF), a naturally occurring riboflavin analogue, has not yet been tested. RoF was first isolated from the soil-dwelling bacterium *Streptomyces davawensis* (26) which recently was described as a valid species and renamed “*Streptomyces davaonensis*” (27). It has been reported that RoF has bactericidal activity against Gram-positive bacteria (26). Within these bacteria, RoF is phosphorylated by the bacterial flavokinase into roseoflavin mononuceotide (RoFMN) and then adenylylated by FAD synthetase into roseoflavin adenine dinucleotide (RoFAD) (28). These flavin cofactor analogs have different physicochemical properties when compared to FMN and FAD. When RoFMN and RoFAD combine with flavoenzymes, they may be rendered inactive (29-32). Another key riboflavin analogue with antimicrobial activity against both Gram-positive and Gram-negative bacteria that has not been tested for antiplasmodial activity is 8-demethyl-8-aminoriboflavin (8AF). 8AF is naturally synthesized as an intermediate product during the synthesis of RoF (33-35).

In this study, we investigated the antiplasmodial activity of RoF and 8AF, as well as an additional eight riboflavin analogues. We show that RoF and 8AF have potent *in vitro* antiplasmodial activity against *P. falciparum* that is counteracted by increasing the extracellular riboflavin concentration. We also tested the effect of RoF against *P. vinckei vinckei* in mice and show that RoF significantly inhibits malaria parasite proliferation *in vivo*.

## Results

### *In vitro* antiplasmodial activity of RoF and 8AF

We initially tested the *in vitro* antiplasmodial activity of RoF and 8AF against the 3D7 strain of *P. falciparum*. RoF and 8AF were found to possess antiplasmodial activity, with IC_50_ values of 1.6 ± 0.1 µM and 7 ± 1 µM, (mean ± SEM, N = 3), respectively (**Figure 2**), when the experiment was carried out in the presence of 0.532 µM riboflavin, the concentration present in standard RPMI-1640. The antiplasmodial activity of RoF decreased by 6-fold (P<0.0001, unpaired t-test) while the activity of 8AF decreased by >3.5-fold, when the extracellular riboflavin concentration was increased from 0.532 to 5 µM. Furthermore, their activity increased by a factor of 53 and 3500 (P<0.0001 and = 0.0046, unpaired t-tests), respectively, when the experiment was carried out in riboflavin-free medium (**Figure 2**). These results are consistent with both compounds exerting their effect on parasite proliferation by competitively inhibiting the parasite’s ability to utilise riboflavin. We next determined the IC_50_ values of RoF and 8AF against *P. falciparum* parasites in the presence of 50 nM riboflavin, a physiologically relevant riboflavin concentration within human plasma (36). Both compounds were found to have IC_50_ values of approximately 120 nM (**Figure 2 and Table 1**).

**Table 1:**
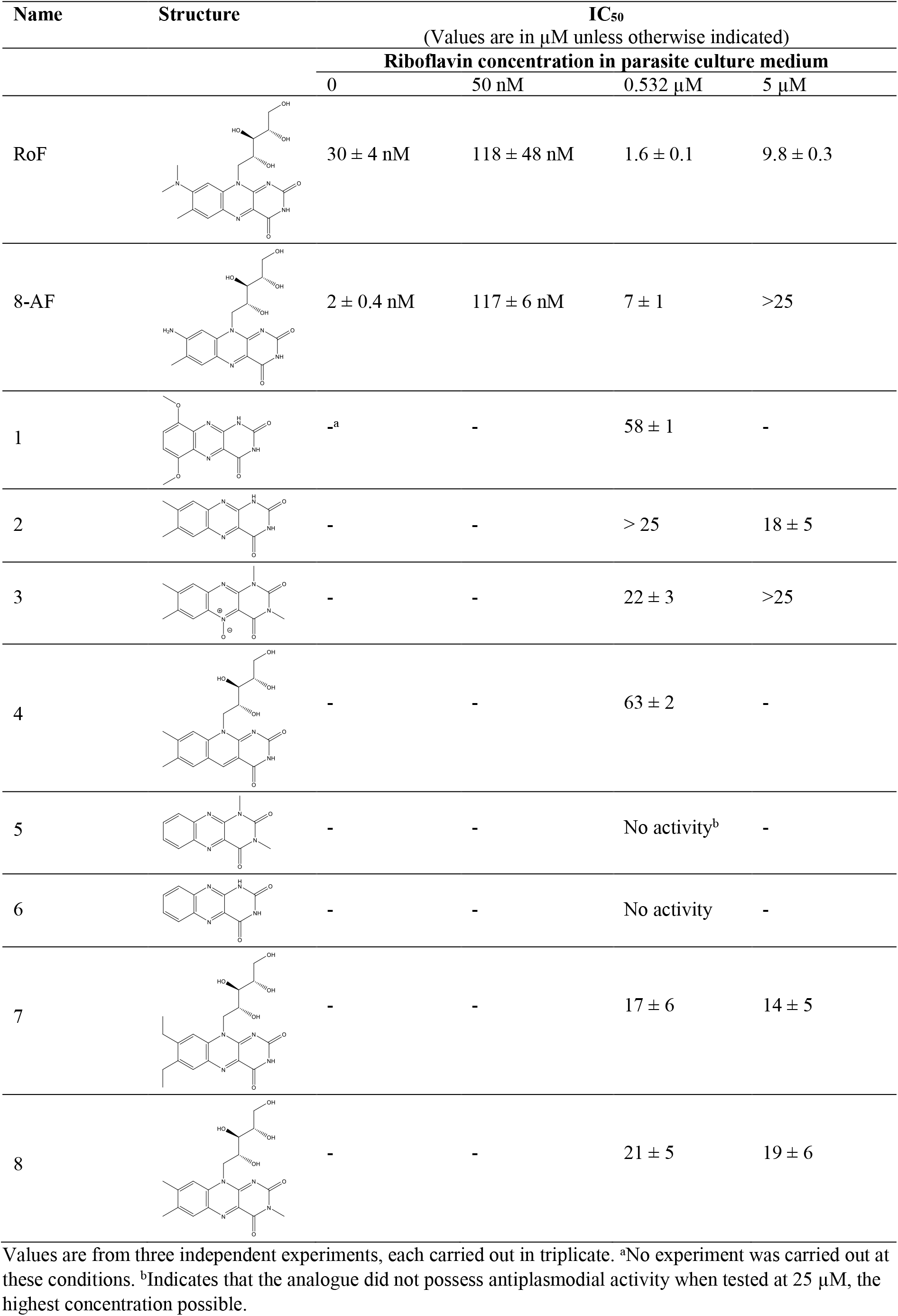
Antiplasomodial activity of riboflavin analogues against *Plasmodium falciparum* in varying riboflavin concentrations.

| Name | Structure | IC<sub>50</sub> |  |  |  |
| --- | --- | --- | --- | --- | --- |
| | | (Values are in $\mu\text{M}$ unless otherwise indicated) | | | |
|  |  | Riboflavin concentration in parasite culture medium |  |  |  |
| | | 0 | 50 nM | 0.532 $\mu\text{M}$ | 5 $\mu\text{M}$ |
| RoF | | $30 \pm 4 \text{ nM}$ | $118 \pm 48 \text{ nM}$ | $1.6 \pm 0.1$ | $9.8 \pm 0.3$ |
| 8-AF | | $2 \pm 0.4 \text{ nM}$ | $117 \pm 6 \text{ nM}$ | $7 \pm 1$ | >25 |
| 1 | | -<sup>a</sup> | - | $58 \pm 1$ | - |
| 2 | | - | - | > 25 | $18 \pm 5$ |
| 3 | | - | - | $22 \pm 3$ | >25 |
| 4 | | - | - | $63 \pm 2$ | - |
| 5 |  | - | - | No activity<sup>b</sup> | - |
| 6 |  | - | - | No activity | - |
| 7 | | - | - | $17 \pm 6$ | $14 \pm 5$ |
| 8 | | - | - | $21 \pm 5$ | $19 \pm 6$ |

Values are from three independent experiments, each carried out in triplicate. <sup>a</sup>No experiment was carried out at these conditions. <sup>b</sup>Indicates that the analogue did not possess antiplasmodial activity when tested at 25 $\mu\text{M}$ , the highest concentration possible.

**Figure 2:**
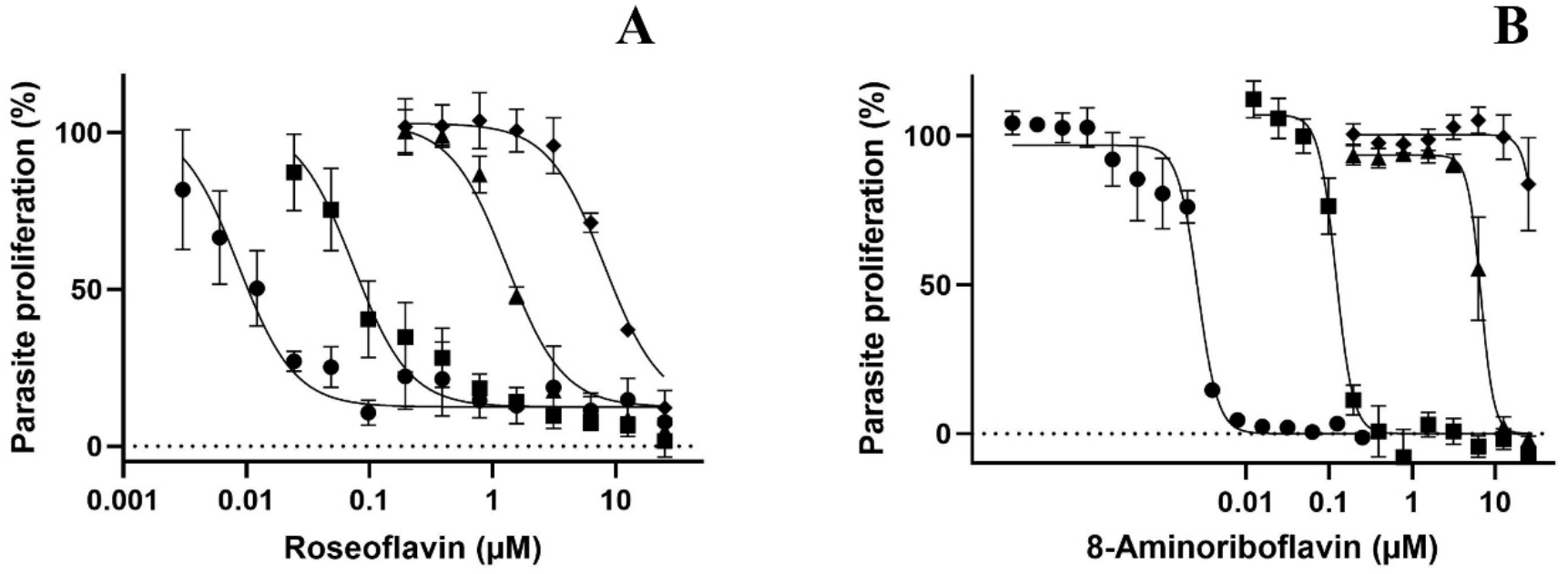
Antiplasmodial activity of RoF (A) and 8AF (B) against *P. falciparum* measured in riboflavin-free medium (circles), or in medium containing 50 nM (squares), 0.532 µM (triangles) or 5 µM riboflavin (diamonds). Values are from three independent experiments, each carried out in triplicate. Error bars represent SEM and, where not shown, are smaller than the symbols.

### *In vitro* antiplasmodial activity of additional riboflavin analogues

Encouraged by the fact that RoF and 8AF were found to possess antiplasmodial activity and that they interfere with riboflavin utilisation, we tested an additional eight riboflavin analogues for activity against *P. falciparum*. Except for compounds **5** and **6**, all the additional analogues were found to possess antiplasmodial activity in RPMI-1640 medium containing 0.532 µM riboflavin. However, their potency was considerably lower than RoF and 8AF (**Table 1**). We then tested the activity of compounds **2, 3, 7**, and **8** in medium containing a 10-fold higher concentration (5 µM) of riboflavin and found that the antiplasmodial activity was unaffected (P>0.579; unpaired t-tes, **Figure S1**), consistent with the compounds either inhibiting parasite proliferation in a manner that although on target, is noncompetitive with riboflavin (although this is unlikely given that the compounds are analogues of riboflavin), or by a mechanism unrelated to riboflavin utilisation.

### *In vivo* antimalarial activity of roseoflavin

In light of the potent *in vitro* antiplasmodial activity of RoF and 8AF, it was important to establish whether the compounds are active *in vivo*. We chose to test RoF because it is commercially readily accessible. The activity of RoF was tested against *P. vinckei vinckei-* infected BALB/c mice using the standard four-day suppression test (37). Infected mice were treated orally with a RoF dose of 150 mg/kg/day or intraperitoneally (IP) with 20 mg/kg/day. Control groups of mice were administered with oral (propylene glycol) or IP (DMSO) vehicle controls only. Similar concentrations of propylene glycol (24) and DMSO (38) have previously been shown to have no effect on *P. vinckei vinckei* growth in mice. Mice that received the initial IP dose, were then given the same dose of RoF for three consecutive days and toxicity was not observed (as determined by weight (**Figure S2**), grooming and level of activity). Mice that were administered the higher dose of RoF (150 mg/kg) orally showed signs of toxicity (**Figure S2**) on the third day (i.e. after the second dose) and were not administered any further doses. RoF (or RoF metabolites, which have the same characteristic colour) was still present in the urine of mice given oral RoF four days after the final administration (evidenced by distinctly pink urine). These mice were euthanised by the seventh day post-infection, but, importantly, we could not detect any parasites in their blood at the time of euthanasia.

The parasitemia in mice administered with RoF IP was measured from the day after the final drug treatment. A 98% reduction in parasitemia was observed in mice two days after four days of IP treatment in comparison with control mice (**Figure 3A**; p = 0.029, unpaired t-test). RoF administration also allowed the mice to survive malaria infection for several additional days following completion of the treatment regime (**Figure 3B**). When the parasitemia reached a value higher than 25%, the mice were euthanased. Euthanasia due to high parasitaemia was the only cause of death of the mice in the IP experiment.

**Figure 3:**
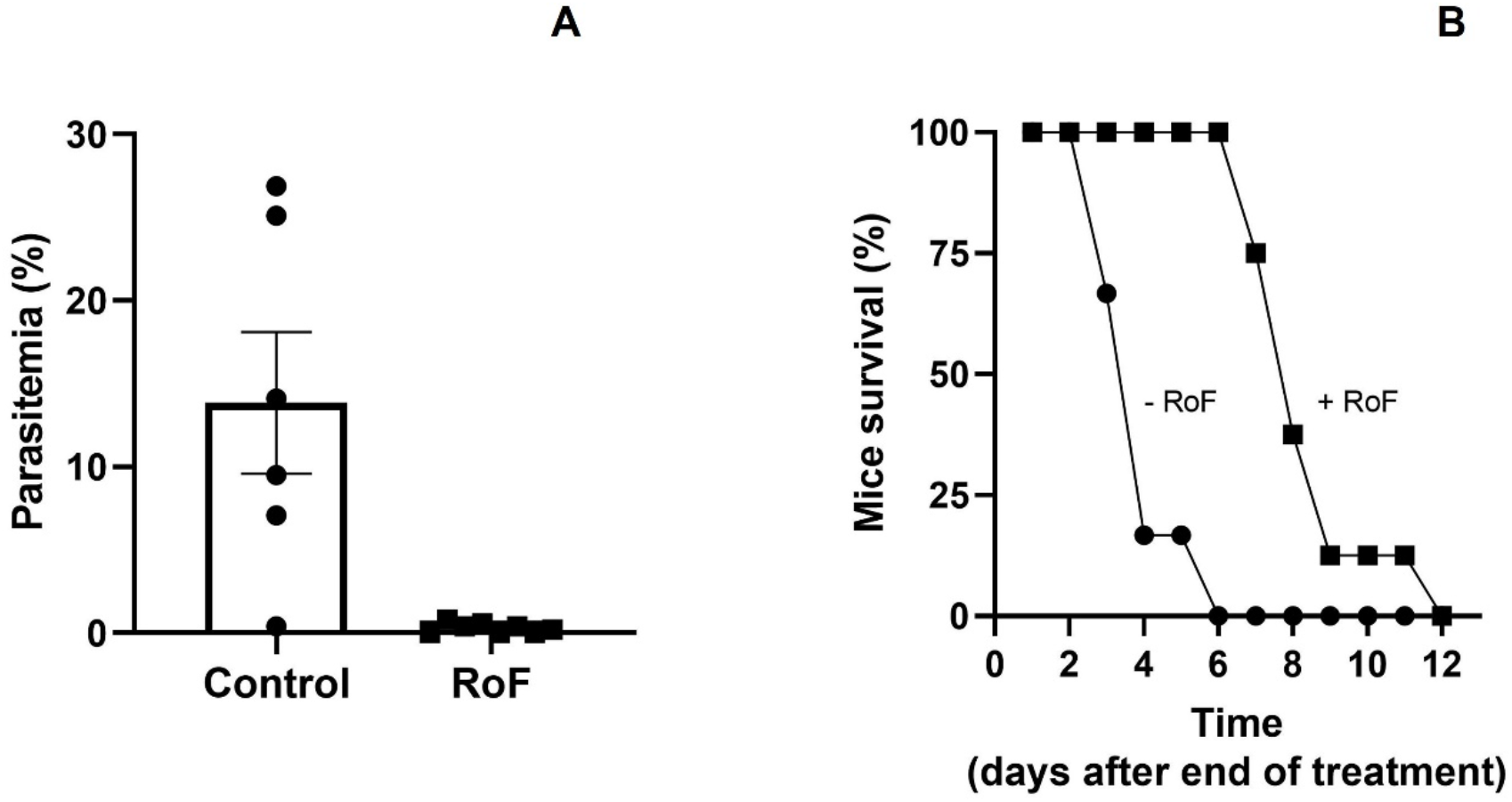
The effect of RoF on the growth of *P. vinckei vinckei in vivo*. (**A**) Average parasitemia in mice determined two days after four days of IP treatment with 20 mg/kg/day RoF or solvent control. Error bars represent SEM. (**B**) Percent mice surviving in the days after completion of the 4-day treatment regime with 20 mg/kg/day RoF (squares) or solvent control (circles).

The relative risk of dying in the control group as compared to that in the RoF group in the ten days following completion of the IP treatment regime was calculated as 8.6 (p < 0.001, from X^2^ = 21.7, log rank test). Hence, IP treatment with 20 mg/kg RoF offered a significant survival advantage to mice infected with *P. vinckei vinckei* compared to the untreated control mice. These findings demonstrate that RoF is effective against malaria parasite proliferation *in vivo*.

## Discussion

The antibacterial activity of RoF and 8AF has been known for some time (26, 31, 39, 40) In this study, we show for the first time that RoF and 8AF kill *P. falcpiarum* parasites at nanomolar concentrations in culture medium containing a riboflavin concentration within the higher end of the human riboflavin plasma levels (2.7 to 42.5 nM, (36)). The observation that the antiplasmodial activity of RoF and 8AF could be altered by changing the extracellular riboflavin concentration is consistent with these compounds targeting riboflavin metabolism, either as inhibitors (reducing the generation of FMN and/or FAD) or substrates of the enzymes involved (thereby generating FMN and/or FAD antimetabolites with a potential to inhibit flavoenzymes). Furthermore, our in *vivo* data demonstrate that RoF (administered at 20 mg/kg/day, IP) significantly reduced the parasitemia and increased the survival time of mice infected with *P. vinckei vinckei*. At this dose, however, the mice were not cured. A 7.5-fold higher dose (150 mg/kg/day) administered orally appeared to completely eliminate the parasites, but was also toxic to the mice. Whilst the oral bioavailability of RoF is encouraging, lower doses will need to be tested to determine which dose/s are efficacious without being toxic.

Riboflavin analogues with electron-donating substituents at position 8 (such as the amino group or alkyl-amino group in the case of 8AF and RoF, respectively) were found to be inert to numerous biological reductants, and hence were assumed incapable of behaving as redox-active molecules (39). These analogues were explored as possible steric replacements for riboflavin, but not as catalytic alternatives (41). As a result, RoFMN, RoFAD and 8AFMN may bind to flavoenzymes that are dependent on FMN or FAD, lowering their activity.

In the last decade, chemical synthesis has been used to create a range of flavin analogues, some of which were found to have strong antibacterial or antiprotist activity (31), but only a few of them have been investigated for potential effectiveness against *P. falciparum*. Encouraged by the potency and on target effects of RoF and 8AF, we tested additional riboflavin analogues. These modifications include riboflavin analogues that lack the ribityl side chain (compound **2**, also known as lumichrome) and it was found to possess off-target activity against *P. falciparum*, but replacing the methyl group of lumichrome at C7 and C8 with hydrogen (compound **6**) and substituting the N1 and N3 of compound **6** with methyl group (compound **5**) led to a complete loss of the activity. However, substituting N5 of compound **5** with the oxide anion (compound **3**) and substituting C6 and C9 of compound **6** with methoxy group (compound **1**) restored the activity. We also found that substituting the N3 of riboflavin with a methyl group (compound **7**), replacing the methyl group at C7 and C8 of riboflavin with ethyl group (compound **8**) and replacing N5 of riboflavin with C atom (compound **4**) resulted in some activity, but they were off-target as determined by the fact that the activity could not be shifted by increasing the extracellular riboflavin concentration (**Figure S1**). Although the results with the additional analogues was disappointing, the results with RoF and 8AF are encouraging and leaves open the possibility that other, more potent and on-target riboflavin analogues could be identified.

## Methods

### *In vitro* cuture of *P. falciparum*

The 3D7 strain of *P. falciparum* was used in all *in vitro* investigations. The parasites were kept in synchronous continuous cultures, as previously described (42). Briefly, *P. falciparum* parasites were maintained in commercial RPMI-1640 medium (Life Technologies) supplemented with 11 mM glucose (Sigma), 24 µg/mL gentamycin (Life Technologies), 200 µM hypothanthine (Sigma) and 0.6% w/v Albumax II (Life Technologies, dissolved in water to 20% (w/v), filter sterilized and stored at (−20°C). Parasites were maintained at 4% hematocrit (HCT), typically in O^+^ erythroyctes in 75 cm^2^ Nunc culture flask, flushed with a gas mixture of 3% CO_2_, 1% O_2_ and 96% N_2_, and held at 37ºC in a horizontal shaking incubator.

The suspension was centrifuged, every 24 hours, at 500 g for 5 min and the supernatant was replaced with fresh medium. The infected erythrocyte pellets were diluted 10-20 times with uninfected erythrocytes when the parasites were in the trophozoite stage, and the parasitemia was not allowed to exceed 5%.

### *In vitro* antiplasmodial activity

#### SYBR-safe assay

The antiplasmodial activity of compounds **1-6** was evaluated using the fluorescence-based SYBR-safe assay (43, 44). Parasites were incubtaed in 96 well plates with riboflavin analogues (compounds **1-6** were obtained from the the National Cancer Institute, DCTD/DTP/DSCB, Rockville, MD, USA, and compound **7** and **8** were purchased from Sigma). All compounds were prepared in DMSO except compound **2** which was dissolved in a mixture of DMSO and 1N KOH. The compounds were tested at a final concentration of 25 µM, except compounds **1, 4 and 8** which were tested at 100 µM. The experiments were carried out at a parasitemia of 0.5 % and a HCT of 1 % for 96 h, starting with parasites in the ring stage. Parasites were incubated with compounds that had been serially diluted (in 2-fold increments). Chloroquine was used as a positive control at a final concentration of 0.5 µM and the corrsponding fluorescence was subtracted from all other values as a background measurment. Drug-free wells were used to represent 100% parasite proliferation. The final volume in each well of the plate was 200 µL. The outermost wells were not used to avoid the “edge effect” (45), but were filled with 200 µL medium. After the 96-hour incubation, the plates were stored at -20°C for at least 24 hours. The plate was then thawed, the contents of the wells resuspended by pipetting and 100 µL transferred to a new plate. To each well, 100 µL SYBR-safe solution in lysis buffer was then added. This solution was made up by adding 2 µL of SYBR-safe stock (Life Technologies) to 10 mL lysis buffer comprised of 5 mM EDTA (Sigma), 20 mM TrisHCl (Sigma), 0.008% w/v saponin (Sigma) and 0.08% v/v Triton X-100 (Sigma). The fluorescence in each well was measured at an excitation of 490 nm and emission of 520nm using a Fluostar Optima fluorometer.

#### Malstat assay

The antiplasmodial activity of RoF, 8AF, **7** and **8** was tested using the Malstat assay because their fluorescent properties interfered with the SYBR-safe assay. This method was carried out according to (46), with minor modifications. Two solutions were prepared to determine the activity of *P. falciparum* lactate dehydrogenase (*Pf*LDH). The first, termed malstat solution, was prepared by dissolving 4 g of sodium L-lactate (Sigma), 1.32 g of Tris (tris(hydroxymethy)aminomethane, Sigma), 22 mg of 3-acetylpyridine adenine dinucleotide (APAD, Sigma) and 0.08% v/v Triton X-100, pH 9 into 50 mL water. This soultion was filter sterilised and stored at 4 °C. The second solution was prepared by mixing 80 mg nitroblue tetrazolium (NBT) and 4 mg phenazine ethosulfate (PES) in 50 mL water. This solution is sensitive to light and was therefore stored in the dark at 4 °C. The antiplasmodial Malstat assay was carried out as described for the SYBR-safe assay up to and including the freezing of the plates. After thawing the plates, the well contents were resuspended and 20 µL of each well transferred to a new plate. To each well 100 µL of the malstat solution was then added, followed by 10 µL of the NBT/PES solution. The plate was incubated in the dark for 45 minutes and the absorbance in each well measured at 620 nm.

### *In vivo* antiplasmodial activity

Approval was obtained from the Australian National University Animal Experimentation Ethics Committee for *in vivo* experiments (approval number F.BMB.31.07). The *in vivo* antiplasmodial activity of RoF was determined *via* a standard four-day suppression test (37). This method assesses the ability of a compound to suppress parasite proliferation when administered in four daily doses. Eight-week-old female BALB/c mice weighing 17-21 g were used. Cryopreserved *P. vinckei vinckei*-infected erythrocytes from a donor mouse were thawed and loaded into a syringe with a 25 G needle. Approximately 3 × 10^6^ erythrocytes (∼1 × 10^6^ of them infected with parasites) in 200 *μ*L of cell suspension were then injected intraperitoneally (IP) into two donor mice. Blood from one of the donor mice was then collected by day six and diluted in saline to 10^7^ infected erythrocytes per 200 μL of cell suspension. A 25 G needle and syringe was used to inject 200 μL of this suspension into each mouse to be used in the four-day suppression test.

Two hours following infection, a group of mice was administered with a 150 mg/kg dose of RoF (37 mM uniform suspension in propylene glycol) orally by gavage. The volume of drug solution given orally to each mouse was approximately 200 μL. Control infected mice were administered with equivalent volumes of propylene glycol to serve as oral vehicle controls. A group of mice was administered IP with a RoF dose of 20 mg/kg (25 mM in DMSO). The volume of drug solution given to each mouse IP was approximately 40 μL. Control mice were administered with equivalent amounts of DMSO to serve as IP vehicle controls. Mice in the IP RoF and corresponding vehicle control groups were given three additional doses approximately 24, 48 and 72 h after the initial dose. Mice in the oral RoF group began to exhibit signs of toxicity after the first two doses. No additional doses were therefore administered.

Blood was taken from the tail (via needle prick) of each mouse 48 h after the final drug administration (72 h after the final oral RoF administration) and used to prepare methanol-fixed, Giemsa-stained smear slides. Microscopic examination was used to determine the parasitemia for each mouse by counting the number of parasitised cells in a random sample of more than 500 erythrocytes. The counting was carried out in a ‘blinded’ fashion and the groups to which the slides belonged to only revealed after all the counting has been completed. Parasitemias were determined daily (except for mice administered oral RoF), and mice with parasitemia >25% were euthanised. Mouse weights were monitored daily from the first drug administration. Mice were observed regularly for signs of toxicity (e.g. loss of weight, lethargy and lack of grooming).

### Statistical analysis

GraphPad Prism 9 was used to do statistical analysis of means using unpaired, two-tailed Student’s t tests. Mouse survival was analysed using the log rank test - a method for determining if two or more independent groups have the same chance of survival. The test compares each group’s whole survival experience and can be considered as an assessment of whether or not the survival curves are equivalent (matching). The relative risk to die in one group, *a*, compared to that in the other group, *b*, is calculated as

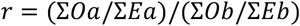

where *O* is the observed number of dead mice, and *E* is the expected number of dead mice. On any one day, *E* is calculated as *E*_*a*_ = (*r*_*a*_ × *d*_*total*_)/*r*_*total*_ where r_a_ is the number of subjects at risk in group *a, d*_*total*_ is the total number of subjects dead from both groups, and *r*_*total*_ is the total number of subjects at risk from both groups.

## Acknowledgements

We are grateful to Dr Kylie Easton for carrying out some of the experiments and to the Canberra Branch of the Australian Red Cross Lifeblood for the provision of red blood cells. AH was supported by a Research Training Program scholarship from the Australian Government and by the Alliance Berlin Canberra ‘‘Crossing Boundaries: Molecular Interactions in Malaria,’’ a program co-funded by the Deutsche Forschungsgemeinschaft (DFG) for the International Research Training Group (IRTG) 2290 and the Australian National University.

## Supplementary Figures

**Figure S1:**
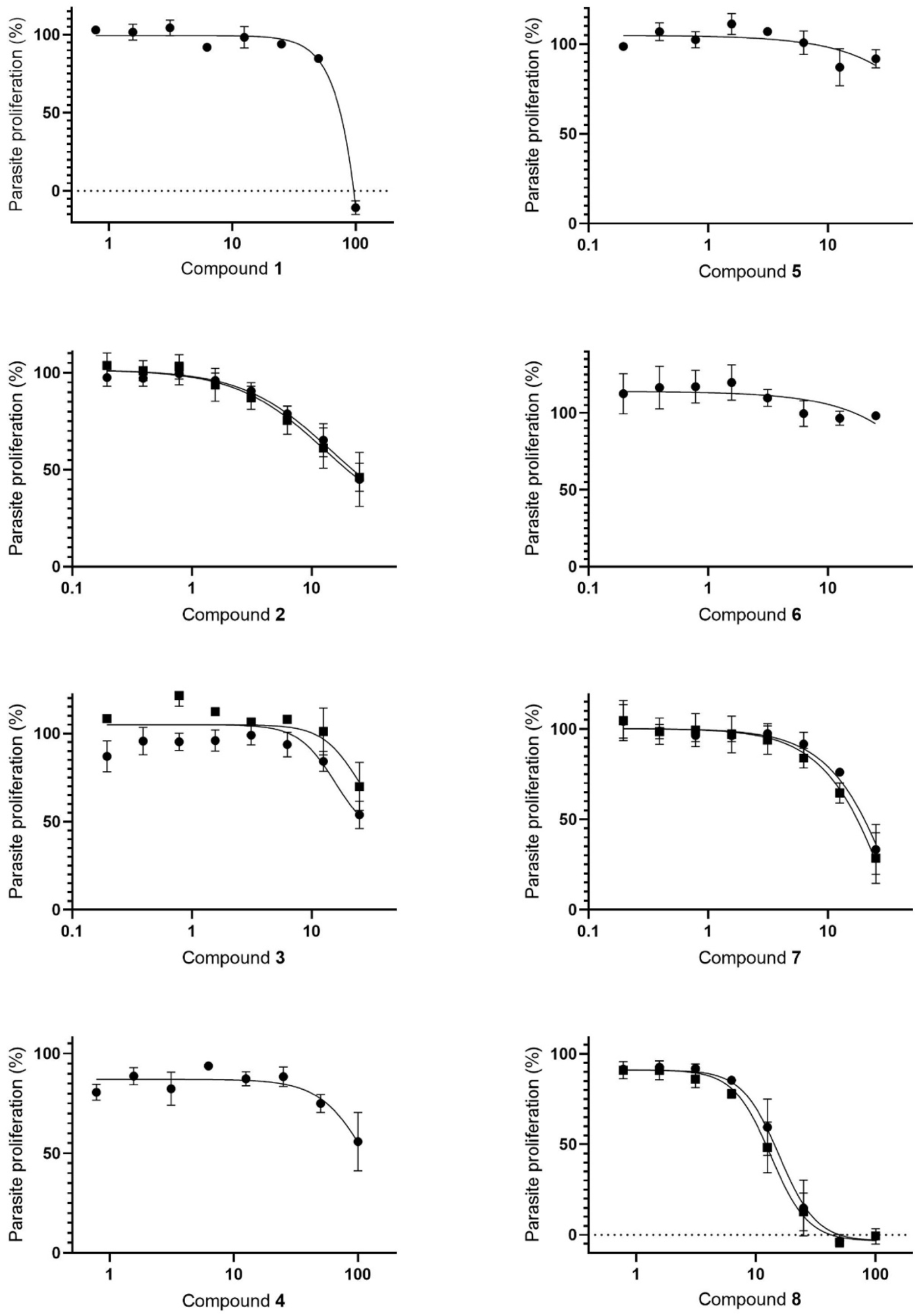
The effect of eight riboflavin analogues on *P. falciparum* proliferation in RPMI-1640 medium containing 0.532 µM riboflavin (circles) or, for compounds 2, 3, 7 and 8, 5 µM riboflavin (squares). Values are averaged from three independent experiments, each carried out in triplicate. Error bars represent SEM.

**Figure S2:**
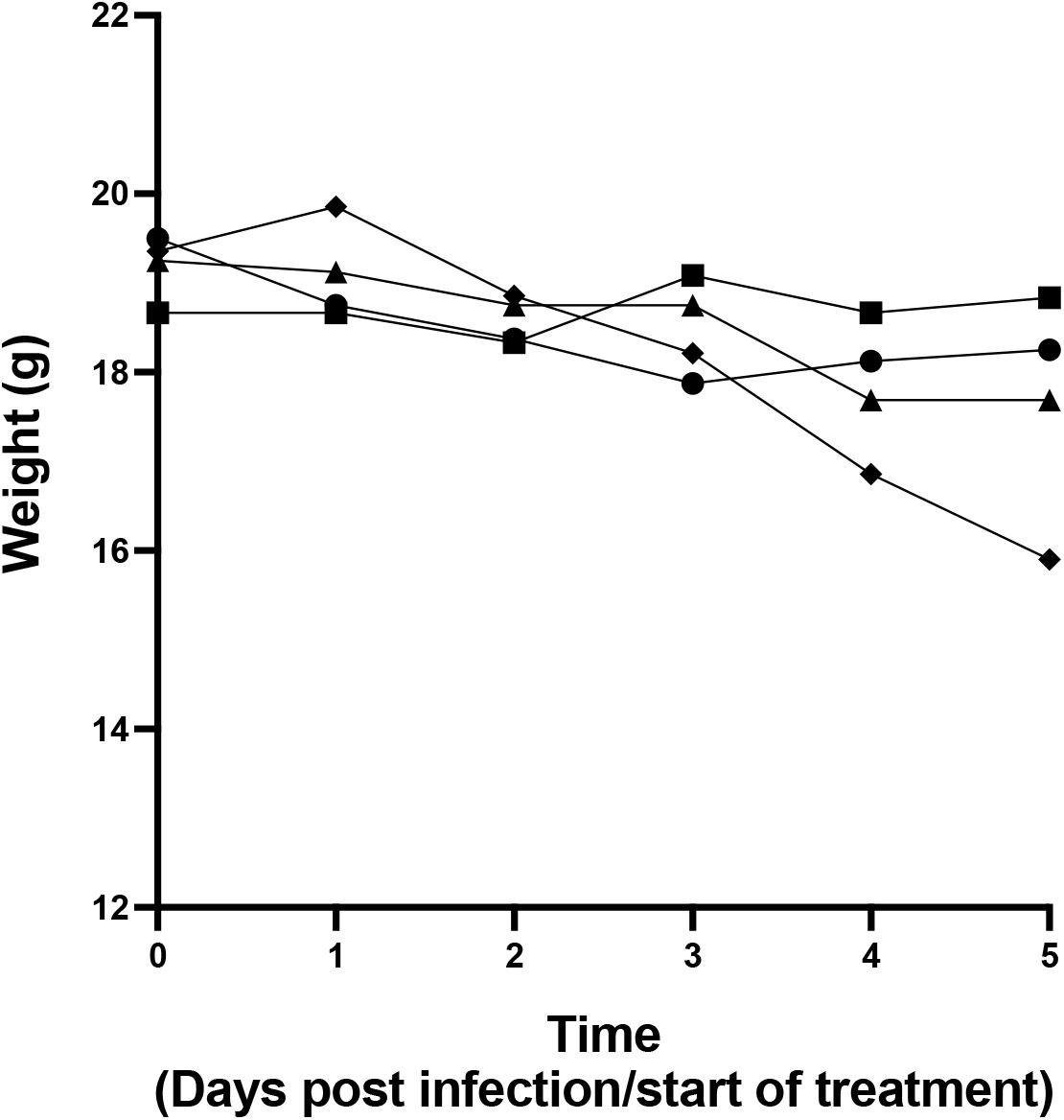
Average weights of mice in each treatment group. Average weight of mice treated with oral vehicle control (circles; n = 4), 150 mg/kg oral roseoflavin (diamonds; n = 7), IP vehicle control (squares; n = 6) and 20 mg/kg IP RoF (triangles; n = 8), over time, starting on the day of infection/start of treatment. Error bars have been omitted for clarity.

